# Inferring generation-interval distributions from contact-tracing data

**DOI:** 10.1101/683326

**Authors:** Sang Woo Park, David Champredon, Jonathan Dushoff

## Abstract

Generation intervals, defined as the time between when an individual is infected and when that individual infects another person, link two key quantities that describe an epidemic: the reproductive number, 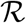, and the rate of exponential growth, *r*. Generation intervals are often measured through contact tracing by identifying who infected whom. We study how observed intervals differ from “intrinsic” intervals that could be estimated by tracing individual-level infectiousness, and identify both spatial and temporal effects, including censoring (due to observation time), and the effects of susceptible depletion at various spatial scales. Early in an epidemic, we expect the variation in the observed generation intervals to be mainly driven by the censoring and the population structure near the source of disease spread; therefore, we predict that correcting observed intervals for the effect of temporal censoring but *not* for spatial effects will provide a spatially informed “effective” generation-interval distribution, which will correctly link *r* and 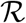. We develop and test statistical methods for temporal corrections of generation intervals, and confirm our prediction using individual-based simulations on an empirical network.

## 1 Introduction

An epidemic can be characterized by the exponential growth rate, *r*, and the reproductive number, 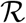. The reproductive number, defined as the average number of secondary cases arising from a primary case, is of particular interest as it provides information about the final size of an epidemic (1; 2). However, estimating the reproductive number directly from disease natural history requires detailed knowledge which is not often available, particularly early in an outbreak (3). Instead, the reproductive number can be *indirectly* estimated from the exponential growth rate, which can be estimated from incidence data (4; 5; 6; 7; 8). These two quantities are linked by generation-interval distributions (9; 10; 11; 12; 13).

A generation interval is defined as the time between when a person becomes infected and when that person infects another person (10). Due to individual variation in infection time, the observed generation-interval distribution can change depending on when and how it is measured (10; 14; 15; 16). There are important distinctions to be made when estimating generation intervals: *intrinsic* generation intervals measure the infectiousness of an infected individual, while *observed* generation intervals refer to the time between actual infection events.

While the intrinsic generation-interval distribution is often assumed to be fixed, the shape of the observed generation-interval distribution depends on when and how intervals are measured (14; 15; 17; 16; 18). When an epidemic is growing exponentially, as often occurs near the beginning of an outbreak, the number of newly infected individuals will be large relative to the number infected earlier on. A susceptible individual is thus relatively more likely to be infected by a newly infected individual. Thus, “backward” generation intervals (which look at a cohort of individuals, and ask when their infectors were infected) will be shorter on average than intrinsic generation intervals – the converse is true when an epidemic is subsiding (15; 16; 18).

In practice, generation intervals are often measured by aggregating available information from contact tracing. While an epidemic is ongoing, these measurements are “censored”: we don’t know what happens after the observation time. These censored intervals are similar to backward intervals: there is a bias towards observing shorter intervals, which are more likely to have concluded by the observation time.

Realized generation intervals are also affected by spatial structure. In a population that does not mix homogeneously, susceptibility will tend to decrease more quickly in the neighbourhood of infected individuals than in the general population. This means that contacts made late in an individual’s infection are more likely to be ineffective due to contacts that were made earlier (because the contactee may have been infected already) As a result, realized generation intervals will have shorter mean than the intrinsic generation-interval distribution. This perspective allows us to reinterpret the finding of (19) that, given an intrinsic generation interval and an observed growth rate, the reproductive number on various network structures is always smaller on a network than would be predicted from homogeneous mixing. These observations allow us to make the following prediction: removing the time-censoring bias from observed generation intervals will yield a spatially corrected “effective” distribution that contains information about the population structure and will allow us to correctly infer the reproductive number from the exponential growth rate.

In this study, we explore spatiotemporal variation in generation intervals measured through contact tracing. We extend previous frameworks to study how the censored generation-interval distributions change over time and compare this distribution with the backward generation-interval distribution. We classify spatial effects on generation intervals into three levels (egocentric, local, and global) and discuss how they affect observed generation-interval distributions. Finally, we compare two methods for accounting for temporal bias and test our prediction using simulations.

## 2 Intrinsic generation-interval distributions

Generation-interval distributions are often considered as population averages, but we can distinguish population-level distributions from individual-level distributions (10; 21); making this distinction clear will be particularly useful when we discuss spatial components (see Fig. 1). An individual-level intrinsic infection kernel *k*(*τ*; *a*) describes the rate at which an infected individual with “aspect” *a* makes “effective contacts” (contacts which will cause infection if the contactee is susceptible). Individual aspects may represent variation in the course of infection (e.g., duration of latent and infectious periods) and the level of infectiousness, which can depend both on biological infectiousness as well as contact patterns.

**Figure 1:**
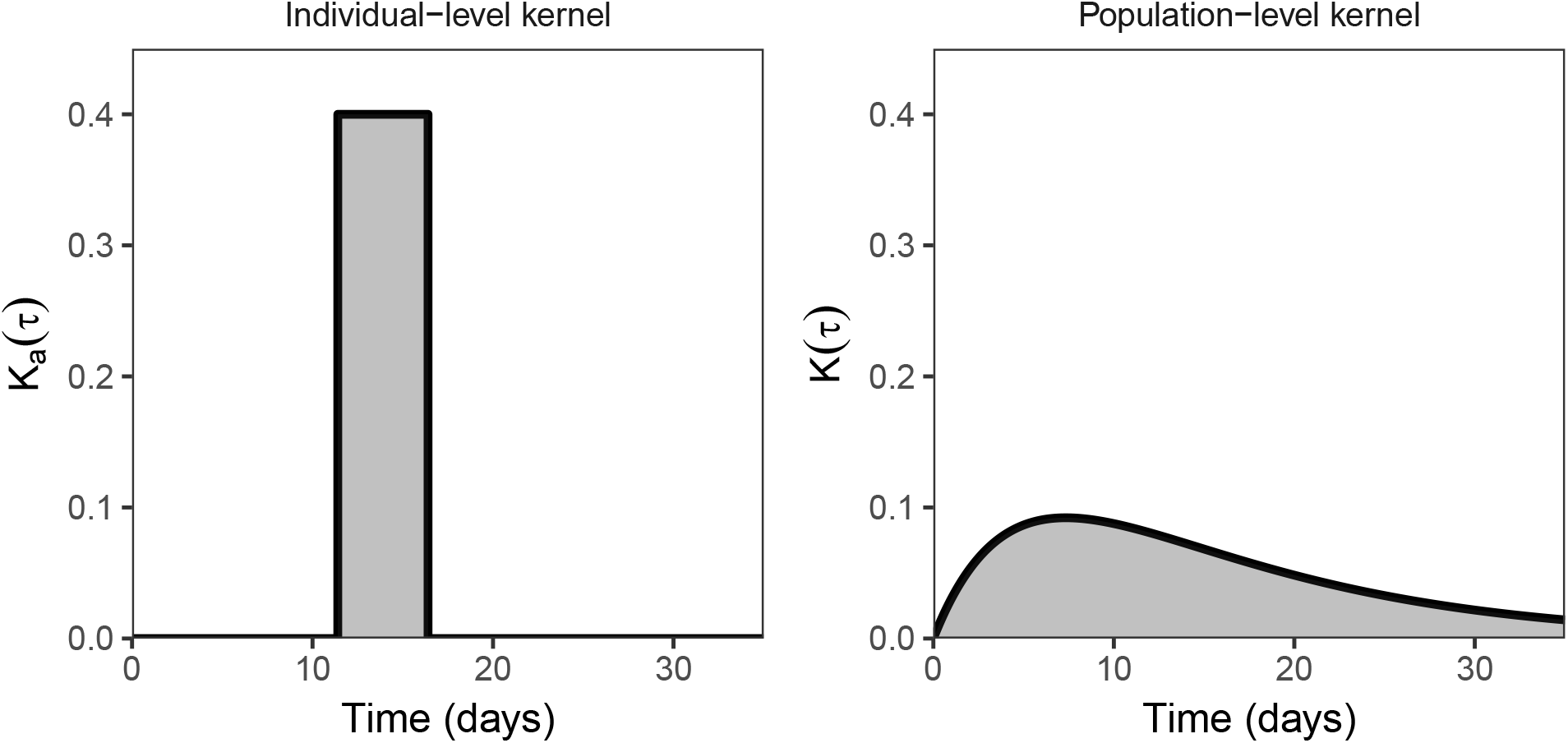
Comparison of individual- and population-level kernels. (Left) an individual-level kernel of an infected individual with latent period of 11.4 days followed by infectious period of 5 days. (Right) a population-level kernel of infected individuals with latent and infectious periods exponentially distributed with means of 11.4 and 5 days, respectively. Shaded areas under the curves are equal to individual- and population-level reproductive numbers, both of which are set to 2 in this example. Parameters are chosen to reflect the West African Ebola outbreak (20).

Assuming that the individual properties are independent of risk of infection, the population-level kernel is given by integrating over these individual variations:

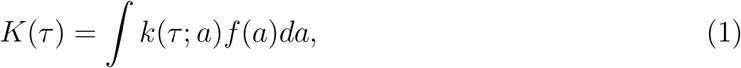

The population-level kernel describes the rate at which secondary infections are expected to be caused by an an infected individual, on *average*. where *f*(*a*) represents a probability density over a (possibly multi-dimensional) aspect space.

Assuming that a population mixes homogeneously, we can write:

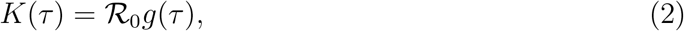

where 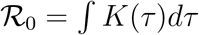 is the basic reproductive number (the expected number of secondary cases caused by a randomly chosen infectious individual in a fully susceptible population), and *g*(*τ*) is the expected time distribution of those cases (the intrinsic generation-interval distribution).

In a homogeneously mixing population, current disease incidence at time *t, i*(*t*), is the product of the current infectiousness of individuals infected in the past and the current proportion of the population susceptible, *S*(*t*).

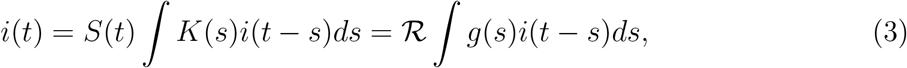

where 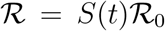 is the effective reproductive number. This model, referred to as the renewal equation, can describe a wide range of epidemic models (22; 23; 24; 25; 12; 11). Over a period of time where the susceptible proportion *S* remains roughly constant, we would expect approximately exponential growth in incidence *i*(*t*); assuming *i*(*t*) = *i*(0)exp(*rt*) yields the Euler-Lotka equation (26), which provides a direct link between speed and strength of an epidemic:

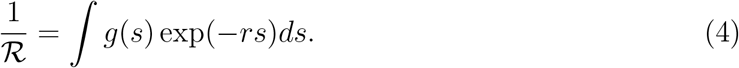

## 3 Generation-interval distributions across time

When generation intervals are estimated through contact tracing during an outbreak, infection events that have not happened yet are not observed. This effect is called “right-censoring”. We can understand the effect of right-censoring using backward generation-interval distributions (17; 15; 16; 18): the observed (censored) generation-interval distribution is a weighted average of backward generation-interval distributions (weighted by incidence) up until the observation time.

The density of new infections occurring at time *t* caused by infectors who were infected at time *t* − *τ* is given by

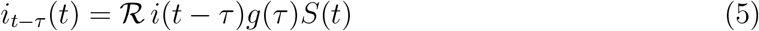

As shown in (16), the backward generation-interval distribution, *b_t_*(*τ*), describes a distribution of infection that occurred *τ* time units before the reference time *t* and is proportional to *i*(*t* − *τ*)*g*(*τ*). The censored generation-interval distribution, *c_t_*(*τ*), describes *all* cohorts infected at time *s* < *t*; thus, the censored generation-interval distribution is just the average of the backward distributions of these cohorts, weighted by incidence:

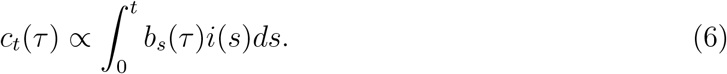

For a single outbreak, the observed mean generation interval through contact tracing will always be shorter than the mean intrinsic generation interval (Figure 2). There are two reasons for this phenomenon. First, longer generation intervals are more likely to be missed due to right censoring (and short generation intervals are more likely to be observed). In particular, when an epidemic is growing exponentially (*i*(*t*) = *i*(0) exp(*rt*)), the initial censored (or backward) generation-interval distribution is just the intrinsic generation-interval distribution discounted by the rate of exponential growth (18):

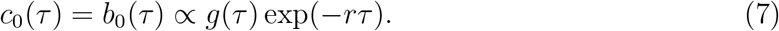

**Figure 2:**
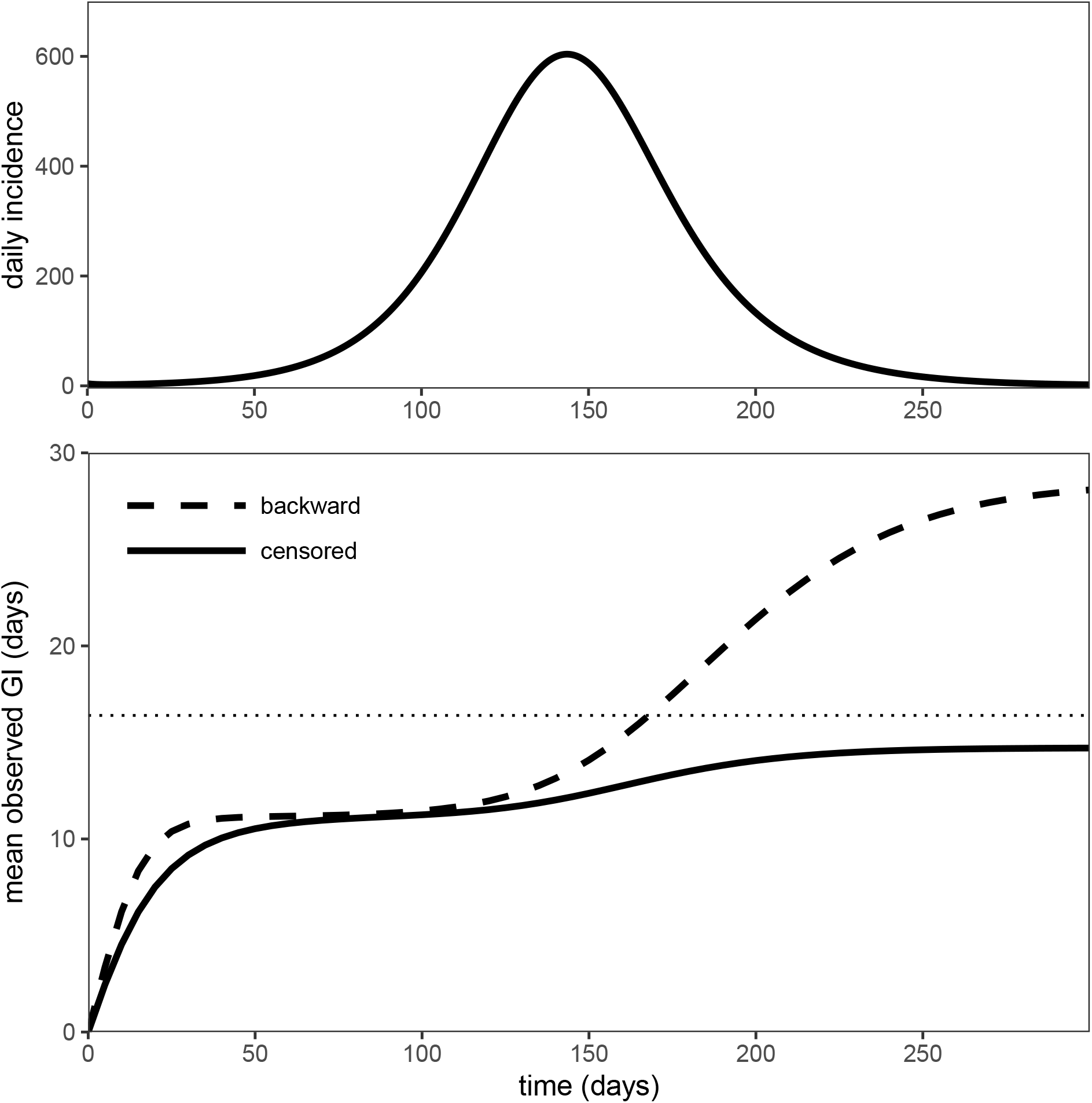
Temporal variation in the mean observed generation interval. A deterministic Susceptible-Exposed-Infectious-Recovered (SEIR) model was simulated using Ebola-like parameters (20): mean latent period 1/*σ* = 5 days, mean infectious period 1/*γ* = 11.4 days, and the basic reproductive number 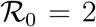. The backward and the censored mean generation interval were calculated over the course of an epidemic. The dotted horizontal line represents the mean intrinsic generation interval.

A deterministic simulation confirms that the censored generation-interval distribution has the same mean as the backward generation-interval distribution during this period (Figure 2). Second, the decreasing number of susceptibles over the course of an epidemic makes long infections less likely to occur (16). Overall, we expect naively using the observed generation-interval distribution, to underestimate the reproductive number.

## 4 Generation-interval distributions across space

Infected individuals may contact the same susceptible individual multiple times, but only the first effective contact gives rise to infection in a given individual (after this, they are no longer susceptible). Therefore, we expect realized generation intervals to be shorter than intrinsic generation intervals, on average, in a limited contact network.

To explore the effect of multiple contacts on realized generation intervals, we first consider the infection process from an “egocentric” point of view, taking into account infectious contacts made by a single infector. We define the egocentric kernel as the rate at which secondary infections are realized by a single primary case with aspect *a* in the absence of other infectors:

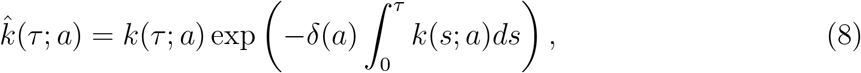

where *k*(*τ*; *a*) is the individual-level intrinsic kernel and 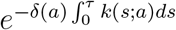 is the probability that a susceptible acquaintance has not yet been contacted by a particular infected individual. The dilution term, *δ*(*a*), models how contacts are distributed among susceptible acquaintances.

Throughout this paper, we assume that there is a constant per-pair contact rate *λ*. In this case, the infectiousness of an individual 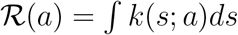 is the product of the number of acquaintances *N*(*a*) and the contact rate *λ*; the dilution term is equal to the reciprocal of the number of acquaintances: *δ*(*a*) = 1/*N*(*a*). This assumption can be relaxed by allowing for asymmetry (19) or heterogeneity (27; 28) in contact rates; for brevity, we do not pursue these directions here.

The population-level egocentric kernel is found by integrating over individual variations:

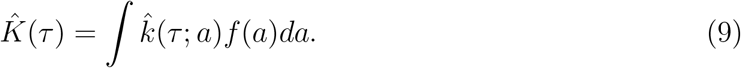

(19) used this same kernel (also assuming a constant per-pair contact rate) to study the effect of network structure on the estimate of reproductive number. The population-level egocentric generation-interval distribution is:

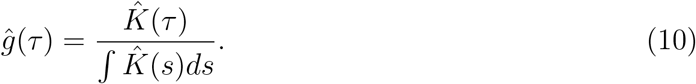

The population-level egocentric generation-interval distribution describes the distribution of times at which secondary infections are realized by an *average* primary case; for convenience, we will often omit “population-level”. Finally, the link between the growth rate and the egocentric reproductive number are linked by the egocentric generation-interval distribution (and the Euler-Lotka equation) (19):

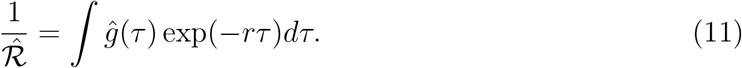

As the egocentric distribution always has a shorter mean than the intrinsic distribution, 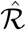 will always be smaller than 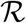 estimated from the intrinsic distribution; this generation-interval-based argument provides a clear biological interpretation for the result presented by (19).

For example, consider a susceptible-exposed-infected-recovered (SEIR) model, which assumes that latent and infectious periods are exponentially distributed. The intrinsic generation-interval distribution that corresponds to this model can be written as (29):

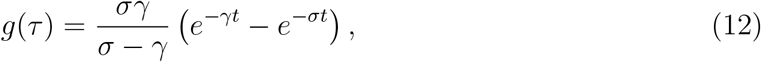

where 1/*σ* and 1/*γ* are mean latent and infectious periods, respectively. Assuming a constant per-pair contact rate of *λ* for any pair, we obtain the following egocentric generation-interval distribution:

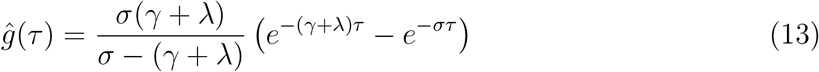

In this case, with fixed infectiousness during the infection period, the effect of accounting for pairwise contacts is the same as an increase in the recovery rate (by the amount of the per-pair contact rate): infecting a susceptible contact is analogous to no longer being infectious (since the contact cannot be infected again); therefore, the resulting egocentric generation-interval distribution is equivalent to the intrinsic generation-interval distribution with mean latent period of 1/*σ* and mean infectious period of 1/(*γ* + *λ*). In practice, directly using this distribution to link *r* and 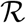 using the Euler-Lotka equation is unrealistic because it requires that we know the per-pair contact rate. Instead, the per-pair contact rate can be inferred from the growth rate *r*, assuming that mean and variance of the degree distribution of a network is known (see (19) supplementary material section 1.4.2); we briefly describe this relationship in Section 7.3.

This calculation can be validated by simulating stochastic infection processes on a “star” network (i.e, a single infected individual at the center connected to multiple susceptible individuals who are not connected with each other). Simulations (Fig. 3) confirm that in this case the distribution of *contact* times matches the intrinsic generation-interval distribution (left panel), while the distribution of realized generation times (i.e., *infection* times) matches the egocentric generation-interval distribution (middle panel).

**Figure 3:**
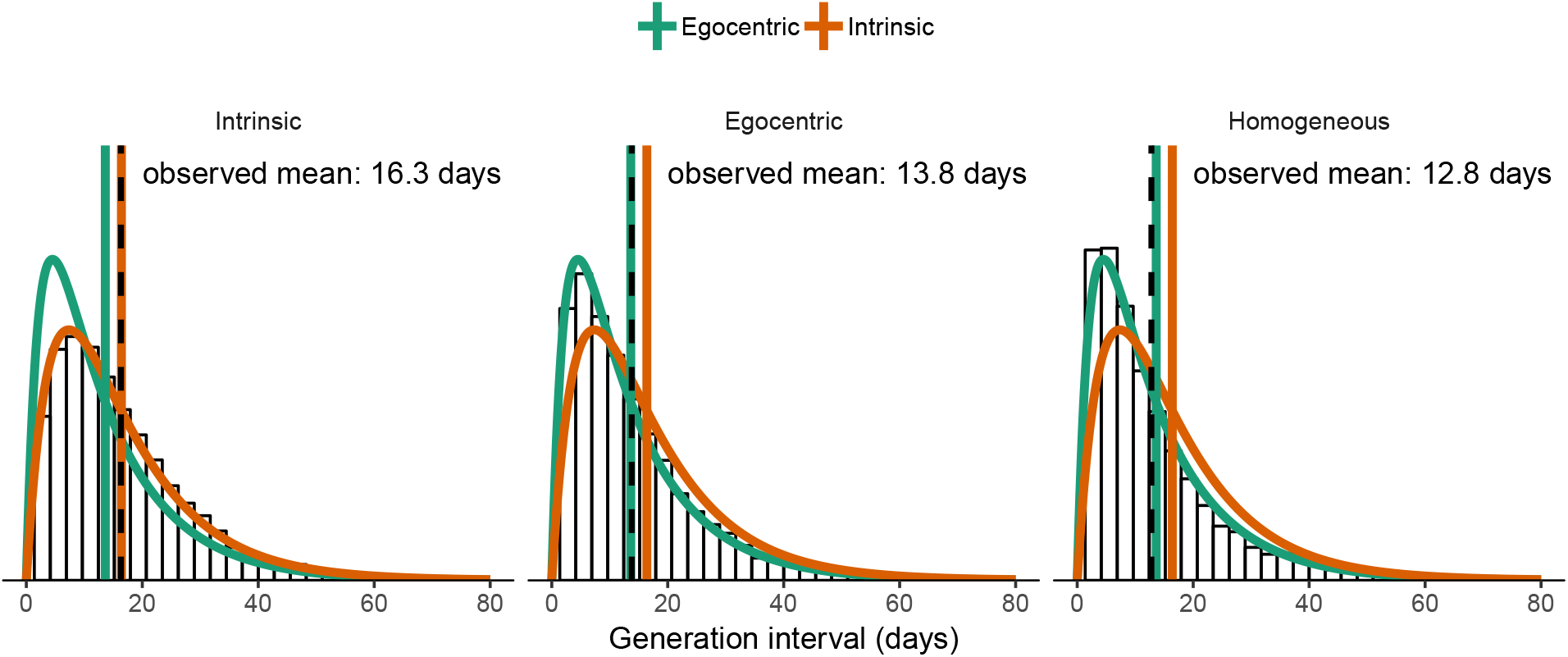
Spatial effects on realized generation intervals. Theoretical distributions and means are shown in color (and are the same in each panel, for reference). Simulated distributions and means are shown in black. (Left) the intrinsic generation-interval distribution corresponds to all contacts by a focal individual, regardless of whether the individual contacted is susceptible. (Middle) the egocentric generation-interval distribution corresponds to the distribution of all infectious contacts by the focal individual with susceptible individuals, in the case where the focal individual is the only possible infector (simulated on a star network). (Right) realized generation-interval distributions are shorter than egocentric distributions in general, because contacts can be wasted when susceptibles become infected through other routes (simulated on a homogeneous network). All figures were generated using 5000 stochastic simulations on a network with 5 nodes (1 infector and 4 susceptibles) with Ebola-like parameters (20): mean latent period 1/*σ* = 5 days and mean infectious period 1/*γ* = 11.4 days. Per-pair contact rate *λ* = 0.25 days^−1^ is chosen to be sufficiently high so that the differences between generation-interval distributions are clear. Each simulation is run until all individuals are either susceptible or has recovered.

The egocentric generation interval (10) only explains some of the reduction in generation times that occurs on most networks, however. Generation intervals are also shortened by indirect connections: a susceptible individual can be infected through another route before the focal individual makes infectious contacts. Simulations on a small homogeneous network confirm this additional effect (Fig. 3, right panel).

In general, spatial reduction in the mean generation interval can be viewed as an effect of susceptible depletion and can be further classified into three levels: egocentric, local, and global. Egocentric depletion, as discussed previously, is caused by an infected individual making multiple contacts to the same individual. Local depletion refers to a depletion of susceptible individuals in a household or neighborhood; we can think of these structures as small homogeneous networks embedded in a larger population structure (and therefore we can expect similar effects as we see in Fig. 3, right panel). Both the egocentric and local depletion effects can be observed early in an epidemic, especially in a highly structured population, even if most of the population remains susceptible. Finally, global depletion refers to overall depeletion of susceptibility at the population level, and explains the reduction in realized compared to intrinsic generation intervals that occurs even in a well-mixed population (Fig. 2).

## 5 Inferring generation-interval distributions from contact-tracing data

When generation intervals are sampled through contact tracing, there will be four effects present in the sample: (1) right-censoring effect, (2) egocentric depletion effect, (3) local depletion effect and (4) global depletion effect. We can correct explicitly for the egocentric effect and, in the case of exponential growth, the right-censoring effect. While the other two effects are difficult to measure, we can make qualitative predictions about their effects on the realized generation intervals and reproductive numbers: both local and global depletion effects reduce number of infections that occur and shorten generation intervals.

Since the right-censoring effect is a sampling bias, we typically want to correct for it. In contrast, spatial effects have the same effect on how the epidemic spreads as they do on observed generation intervals. We therefore expect that starting from observed generation intervals and correcting for the right-censoring effect, will allow us to estimate an “effective” generation interval that accurately reflects dynamics of spread. When temporal correction is performed early in an outbreak, during the exponential growth phase, the effective distribution should incorporate egocentric and local spatial effects but not the global effects; we expect this distribution to correctly link *r* and 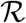. We will call the temporally corrected effective generation-interval distribution *g*_eff_(*τ*). In a large homogeneously mixing population, the effective distribution is expected to be equivalent to the intrinsic distribution (since the egocentric and local depletion are negligible in the absence of network structure, and global depletion is negligible early in the epidemic).

Here, we investigate two methods for correcting for temporal bias in contact-tracing data (see Methods for details). We refer to the first method as the population-level method as it relies on the observed distribution aggregated across the entire population. When an epidemic is growing exponentially, right-censoring causes the observed generation interval to be discounted by the exponential growth rate (7); hence, we can “undo” the censoring by exponentially weighting the observed generation-interval distribution (17; 15; 18):

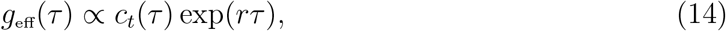

where *r* is the exponential growth rate.

We refer to the second method as the individual-level method because it relies on individual contact information. We model each infection as a non-homogeneous Poisson process arising from the infector (23); incorporating information about time of infection of an infector, time of infection of an infectee, and time since the beginning of an epidemic allows us to explicitly model the censoring process in the observed generation intervals. For both methods, mean and coefficient of variation (CV) of generation-interval distributions are estimated by maximum likelihood; the inferred generation-interval distribution is then used to estimate the reproductive number 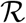 from the observed growth rate *r* using the Euler-Lotka equation.

To test these methods, we simulate 100 epidemics with Ebola-like parameters on an empirical network (30) and compare the estimates of reproductive number with empirical reproductive numbers, which we define as the average number of secondary cases generated by the first 100 infected individuals (Fig. 4). As expected, calculating reproductive number based on the intrinsic generation-interval distribution overestimates the empirical reproductive number; estimates based on the egocentric generation-interval distribution (10) address this problem only partially, as they do not account for indirect (local) spatial effects. Direct estimates based on the observed generation intervals from contact tracing severely under-estimate the empirical estimates. While both population- and individual-level corrections provide similar estimates to the empirical reproductive number on average, population-level estimates are more variable as they are more sensitive to outliers in generation intervals and our estimate of the growth rate. For smaller values of 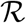, we expect the differences to become smaller. In the electronic supplementary material, we present the same figure using smaller 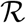 (Appendix A.1) and using Erlang-distributed latent periods (Appendix A.2), which better corresponds to Ebola. Overall, our qualitative conclusions do not change.

**Figure 4:**
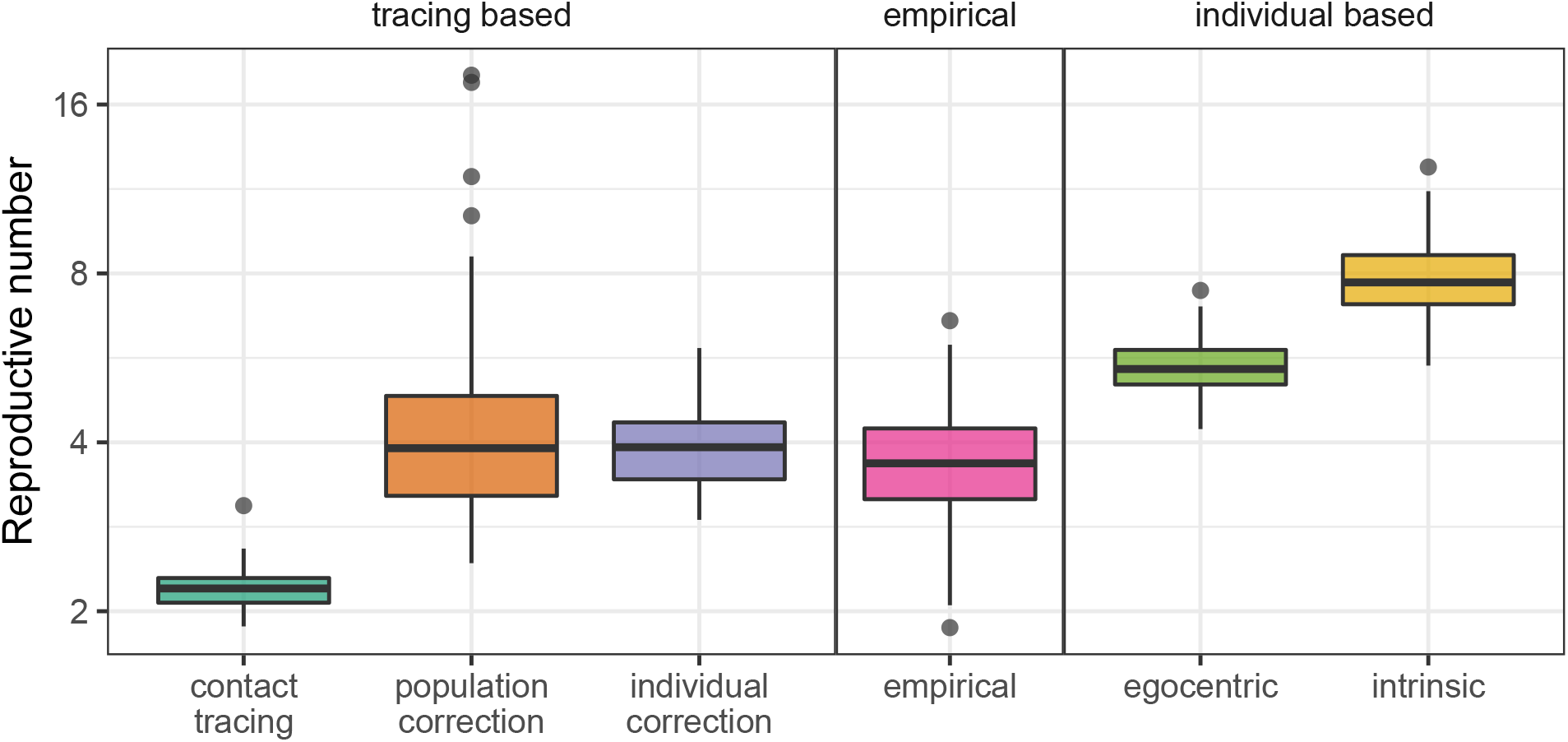
Comparison of estimates of reproductive number based on various methods. Using the observed generation-interval distributions (based on the first 1000 generation intervals) without correcting for right-censoring severely underestimates the reproductive number. Similarly, using the intrinsic generation-interval distribution overestimates the reproductive number because it fails to account for local spatial effects; the egocentric distribution corrects for this only partially. Both population-level and individual-level methods provide estimates of reproductive number that are consistent with the empirical estimates, which we define as the average number of secondary cases generated by the first 100 infected individuals, but the individual-level method is more precise. Boxplots are generated using 100 stochastic simulations of the SEIR model on an empirical network using Ebola-like parameters (20): mean latent period 1/*σ* = 5 days and mean infectious period 1/*γ* = 11.4 days. Per-pair contact rate *λ* = 0.08 days^−1^ is chosen to be sufficiently high such that differences are clear.

## 6 Discussion

The intrinsic generation-interval distribution, which describes the expected time distribution of secondary cases, provides a direct link between speed (exponential growth rate, *r*) and strength (reproductive number, 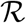) of an epidemic (12; 10; 21; 13). However, observed generation-interval distributions can vary depending on how and when they are measured (15; 17; 16; 18); determining which distribution correctly links *r* and 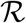 can be challenging. Here, we analyze how observed generation intervals, measured through contact tracing, differ from intrinsic generation intervals. Changes due to temporal censoring reflect observation bias, whereas changes due to spatial or network structure reflect the dynamics of the outbreak. Thus, correcting the observed distribution for temporal, but not spatial, effects provides the correct link between *r* and 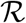.

Observed generation intervals are subject to right-censoring – it is not possible to trace individuals that have not been infected yet. These right-censored distributions can be thought of as averages of “backward” generation intervals (measured by tracing the infectors of a cohort of infected individuals) (14; 15; 17; 16; 18). During an ongoing outbreak, the observed generation-interval distribution will always have a shorter mean than the intrinsic-interval distribution due to right-censoring. Early in the outbreak, censored intervals are expected to match the early-outbreak backward intervals. Near the end of an outbreak, the effect of right-censoring becomes negligible but the observed generation intervals are still shorter on average than intrinsic generation intervals, because of depletion of the susceptible population.

We think of susceptible depletion as operating on three levels: egocentric, local, and global. Egocentric susceptible depletion refers to the effect of an infected individual making multiple contacts to the same susceptible individual. Accounting for the egocentric effect allowed us to link the results by (19) to established results based on generation intervals. Local susceptible depletion refers to the effect of other closely related infected individuals (e.g., in a household or neighborhood) making multiple contacts to the same susceptible individual. Global susceptible depletion refers to the overall decrease in the susceptible pool in the population.

Susceptible depletion happening at all three levels shortens realized generation intervals but acts on different time scales. Egocentric and local depletion effects are present from the beginning of an epidemic, even when depletion in the global susceptible population is negligible, and can strongly affect the initial spread of an epidemic. Therefore, we predicted the observed generation intervals during an exponential growth phase to contain information about the contact structure, allowing us to estimate the *effective* generation-interval distribution by simply accounting for the right-censoring. Simulation studies confirmed our prediction: using the effective generation-interval distribution provides the correct link between *r* and 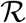.

We compare two methods for estimating the effective generation-interval distribution and assume that the effective generation-interval distribution follows a gamma distribution. The gamma approximation of the generation interval distribution has been widely used due to its simplicity (31; 6; 32; 33; 34); we previously thought that summarizing the entire distribution with two moments (mean and variance) is sufficient to understand the role of generation-interval distributions in linking *r* and 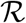 (13). However, further investigation of our methods suggests that making a wrong distributional assumption can lead to biased estimates of the mean and CV of a generation-interval distribution (Appendix A.3), even though the gamma distribution “looks” indistinguishable from the shape of the intrinsic generation-interval distribution (derived from the SEIR model). These results are particularly alarming because it is impossible to know the true shape of the generation-interval distribution for real diseases. Nonetheless, biases in the parameter estimates of a generation-interval distribution may cancel out and give unbiased estimate of the reproductive number (Appendix A.3).

Generation-interval-based approaches to estimating the reproductive number often assume that an epidemic grows exponentially (9; 12; 11; 13). In practice, heterogeneity in population structure can lead to subexponential growth (35; 36; 37; 38; 39; 40); we therefore expect our simulations on an empirical network to be better characterized by subexponential growth models (40). However, our simulations suggest that the initial exponential growth assumption still provides a viable approach for estimating the reproductive number.

Contact tracing provides an effective way of collecting epidemiological data and controlling an outbreak (41; 42; 43). In particular, using tracing information allows us to infer real-time estimates of the time-varying reproductive number (44; 45; 46; 47); generation-interval distributions, which can be either assumed or estimated, often play a central role in analyzing tracing data. Our study illustrates that generation intervals measured through contact-tracing data contain information about the underlying contact structure, which can be implicitly reflected in the estimates of the reproductive number; this perspective can be particularly useful for characterizing an epidemic because detailed information about the contact structure is often unavailable.

The generation-interval distribution is a key, and often under-appreciated, component of disease modeling and forecasting. Different definitions, and different measurement approaches, produce different estimates of these distributions. We have shown that estimates based on observed generation intervals (e.g., through contact tracing) differ in predictable ways from estimates based on underlying measures of infectiousness (e.g., from shedding studies). These predictable differences can arise from temporal effects, egocentric spatial effects, local spatial (or network) effects and population-level effects. We have shown that correcting observed intervals for temporal effects allows us to estimate a spatially informed “effective” distribution, which accurately describes how disease spreads in a population. Future studies should carefully consider how measurement influences estimated generation-interval distributions, and how these distributions influence the spread of disease.

## 7 Methods

### 7.1 Deterministic SEIR model

To study the effects on right-censoring on the observed generation intervals, we use the deterministic Susceptible-Exposed-Infectious-Recovered (SEIR) model. The SEIR model describes how disease spreads in a homogeneously mixing population; it assumes that infeceted individuals become infectious after a latent period. We use a SE^*m*^I^*n*^R model, which extends the SEIR model to have multiple equivalent stages in the latent and infectious periods. This gives latent and infectious periods with Erlang distributions (gamma distributions with integer shape parameters), which are often more realistic than the exponentially distributed periods in the standard SEIR model (48; 49):

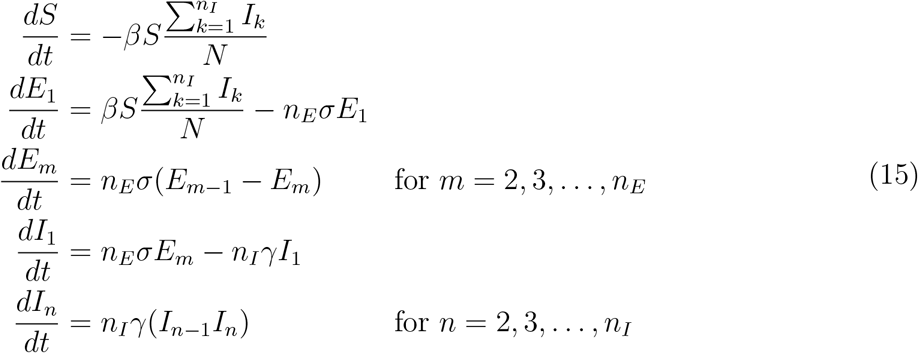

where *S* is the number of susceptible individuals, *E_m_* is the number of exposed individuals in the *m*-th compartment, *I_n_* is the number of infectious individuals in the *n*-th compartment, and *R* is the number of recovered individuals. Parameters of the model are specified as follows: *N* is the total population size, *β* is the transmission rate, 1/*σ* is the mean latent period, *n_E_* is the number of latent compartments, 1/*γ* is the mean infectious period, and *n_I_* is the number of infectious compartments. We show results based on more realistic distributions in Appendix.

### 7.2 Stochastic SEIR model

We simulated an individual-based SEIR model on a contact network, using an algorithm based on the “Gillespie algorithm” (50; 51). We begin by randomly selecting individuals assumed to be infected at *t* = 0. For each infected individual *i*, we randomly draw the latent period *E_i_* from an Erlang distribution with mean 1/*σ* and shape *n_E_*. We then construct the random infectious period and infectious contact times simultaneously as follows. For each of the *n_I_* stages of the infectious period, we draw the number of effective contacts (before transitioning to the next compartment) from a geometric distribution with probability *n_I_γ*/(*S_i_λ* + *n_I_γ*), where *S_i_* is the number of susceptible acquaintances and *λ* is the per-pair contact rate. We then choose the time between consecutive events (the chosen number of contacts, followed by exit from the given stage of infection) from an exponential distribution with rate *S_i_λ* + *n_I_γ*. For each contact, a contactee is uniformly sampled from the set of susceptible acquaintances of the individual *i*. The infectious period *I_i_* is the sum of all of these waiting times.

After repeating the contact process for all initially infected individuals, all contacts are put into a sorted queue. The first person in the queue becomes infected, and the current time is updated to infection time of this individual. Any subsequent contacts made to this individual are removed from the queue because they will no longer be effective. We repeat the contact process for this newly infected individual. Then, new contacts are added to the sorted queue. The simulation continues until there are no more contacts are left in the queue.

### 7.3 Egocentric relationship between *r* and 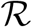 (SEIR model)

Here, we show that the egocentric relationship between *r* and 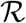 derived by Trapman (19) (see the original source for detailed derivations) matches what would be calculated by applying the Euler-Lotka equation to the egocentric (rather than the intrinsic) generation-interval distribution. Assume that latent and infectious periods are exponentially distributed with mean 1/*σ* and 1/*γ*, respectively. Assuming a constant per-pair contact rate of *λ* for any pair, the egocentric generation-interval distribution can be written:

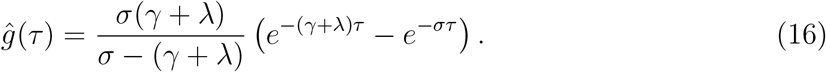

Substituting into (11), we get

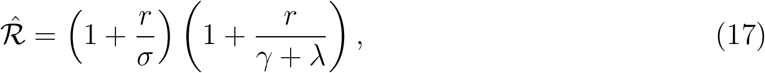

where *r* is the exponential growth rate. Alternatively, the reproductive number can be expressed based on the degree distribution (mean *μ* and variance *σ*^2^) of a network:

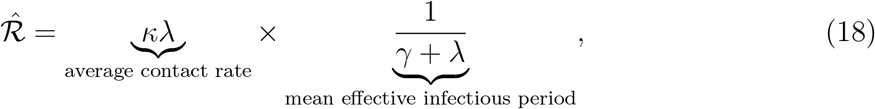

where *κ* = *σ*^2^/*μ* + *μ* − 1, referred to as the mean degree excess (52), describes the expected number of susceptible individuals that an average infected individual will encounter early in an outbreak. Combining the two equations, we get

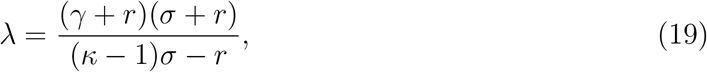

which completes the relationship between the growth rate and the egocentric reproductive number:

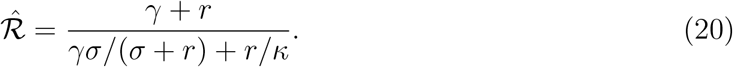

### 7.4 Estimating generation-interval distributions from contact-tracing data

The **Population-level method** estimates the effective generation-interval distribution by reversing the inverse exponential weighting in the observed generation-interval distribution:

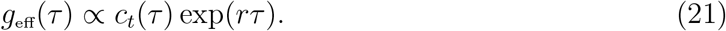

In order to do so, we first fit a gamma distribution to the observed generation-interval distributions; specifically, we estimate the mean *Ḡ* and shape *α* of a gamma distribution by maximum likelihood. Then, the effective generation-interval distribution also follows a gamma distribution with mean *α*/(*α*/*Ḡ* − *r*) and shape *α*.

The **Individual-level method** models each infection *i* from an infected individual *j* as a non-homogeneous Poisson process between the time at which infector *j* was infected (*t_j_*) and the censorship time (*t*_censor_), with time-varying Poisson rate at time t is equal to 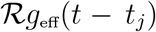, where *g*_eff_(*t*) is the effective generation-interval distribution (53). We use a gamma distribution (parameterized by its mean and shape) to model the effective generation-interval distribution. Then, the probability that an individual *j* infects *n_j_* individuals between *t_j_* and *t*_censor_ is equal to

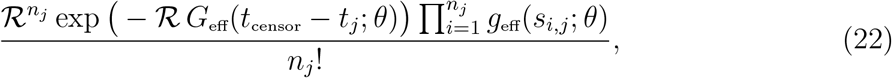

where *s_i,j_* is the observed generation interval between infector *j* and infectee *i*, and *θ* is a (vector) parameter of the generation-interval distribution *g* (and the corresponding cumulative distribution function *G*_eff_).

The likelihood of the non-homogeneous poisson process can be written as:

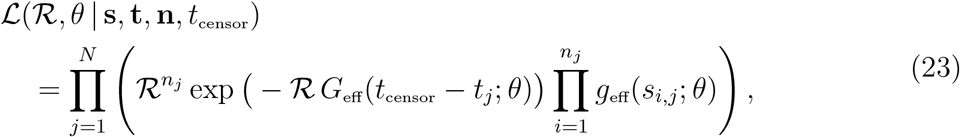

where *N* is the total number of infected individuals. Here, we estimate parameters 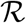 and *θ* by maximum likelihood. Since the estimate of 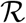 is sensitive to under-reporting, we use the estimated generation-interval distribution to infer the effective reproductive number from the estimated growth rate (using the Euler-Lotka equation). (54) proposed a related approach based on discretized incidence reports.

### 7.5 Measuring the exponential growth rate

We estimate the exponential growth rate *r* of an epidemic from daily incidence by modeling the cumulative incidence *c*(*t*) with a logistic function (8). Fitting directly to cumulative incidence can lead to overly confident results (55); instead we fit interval incidence *x*(*t*) = *c*(*t* + Δ*t*) − *c*(*t*), where Δ*t* is 1 day, to daily incidence, assuming that daily incidence follows a negative binomial distribution. We estimate the logistic parameters *r, K, c*_0_, and *θ* by maximum likelihood. Fitting time window is defined from the last trough before the peak of an epidemic to the first day after the peak of an epidemic.

### 7.6 Empirical network

To simulate epidemics on a realistic network, we use the ‘condensed matter physics’ network from the Stanford Large Network Dataset Collection (30). This graph describes co-authorship among anyone who submitted a paper to Condensed Matter category in the arXiv between January 1993 and April 2003 (56). It consists of 23133 nodes and 93497 edges. The same network was used by (19) to study how network structure affects the estimate of the reproductive number.

## Authors’ contributions

SWP led the literature review, wrote the first draft of the MS, performed analytic calculations and simulations; JD conceived the study and performed analytic calculations; all authors contributed to refining study design, literature review, and final MS writing. All authors gave final approval for publication.

