## Supplementary file for "Inferring generation-interval distributions from contact-tracing data"

### A Appendix

#### A.1 Comparison of estimates of reproductive number – smaller $\mathcal{R}$

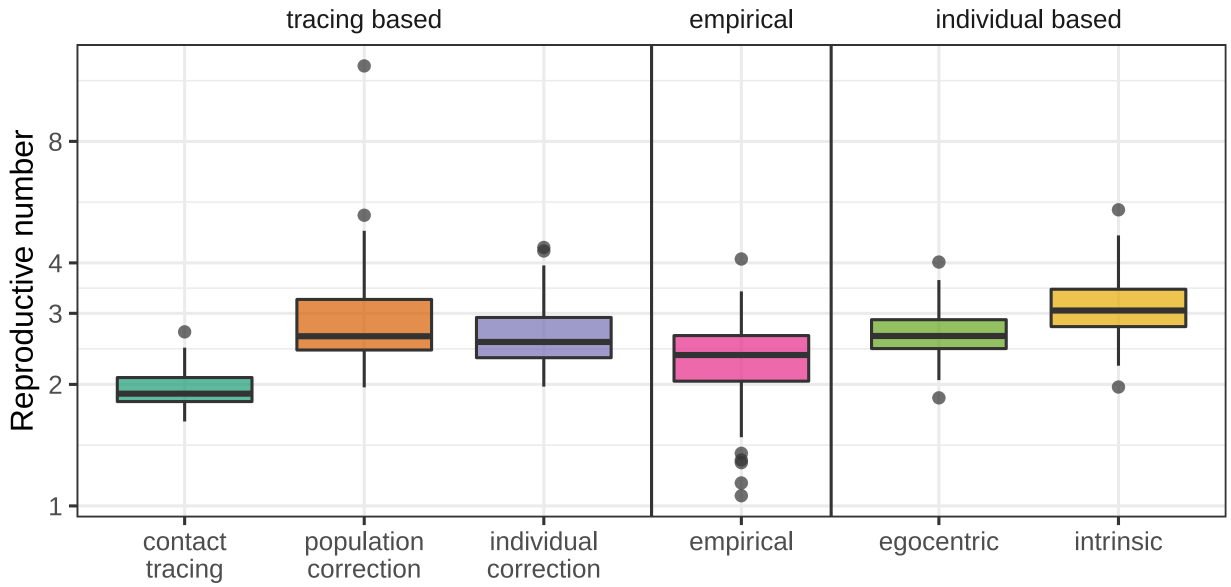


Figure A.1: **Comparison of estimates of reproductive number based on various methods.** This figure matches Fig. 4 in the main text. A smaller per-pair contact rate ($\lambda = 0.0026$ days$^{-1}$) is used to simulate epidemics. All other parameters are the same as in Fig. 4 in the main text.

#### A.2 Comparison of estimates of reproductive number – Erlang distributed latent periods

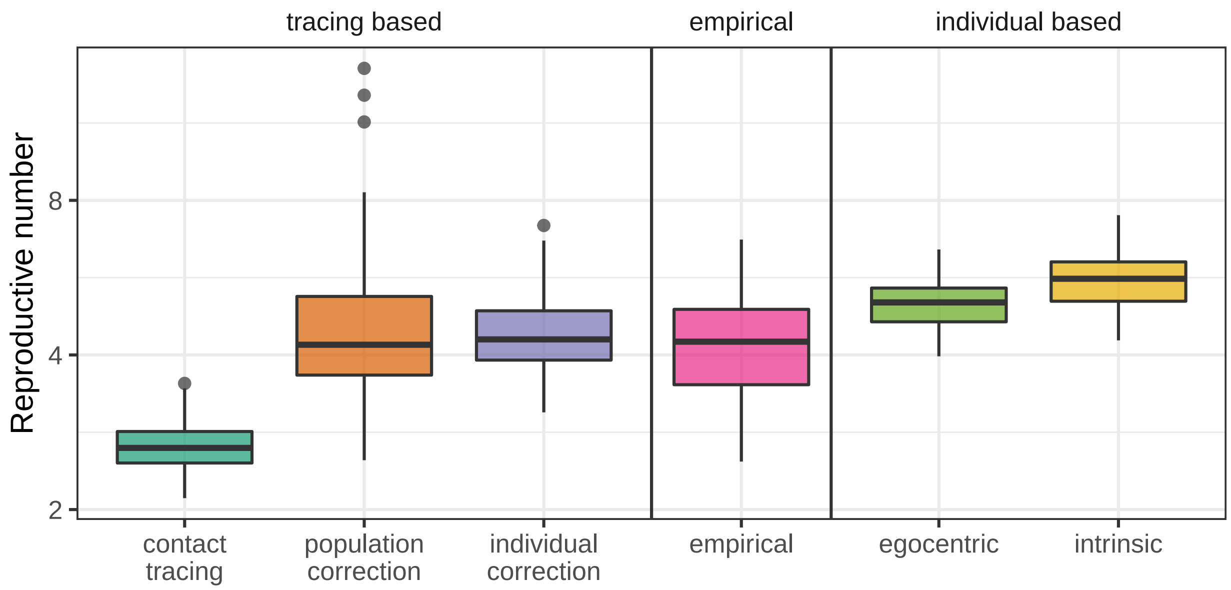


Figure A.2: **Comparison of estimates of reproductive number based on various methods.** This figure matches Fig. 4 in the main text which used a SEIR model. Here, an Erland-distributed latent period ($n_E = 2$) is used to simulate epidemics; this assumption better matches the incubation period distribution of the Ebola virus disease than the exponential assumption (1). All other parameters are the same as in Fig. 4 in the main text.

#### A.3 Testing the individual-based method on simulations on a homogeneous network

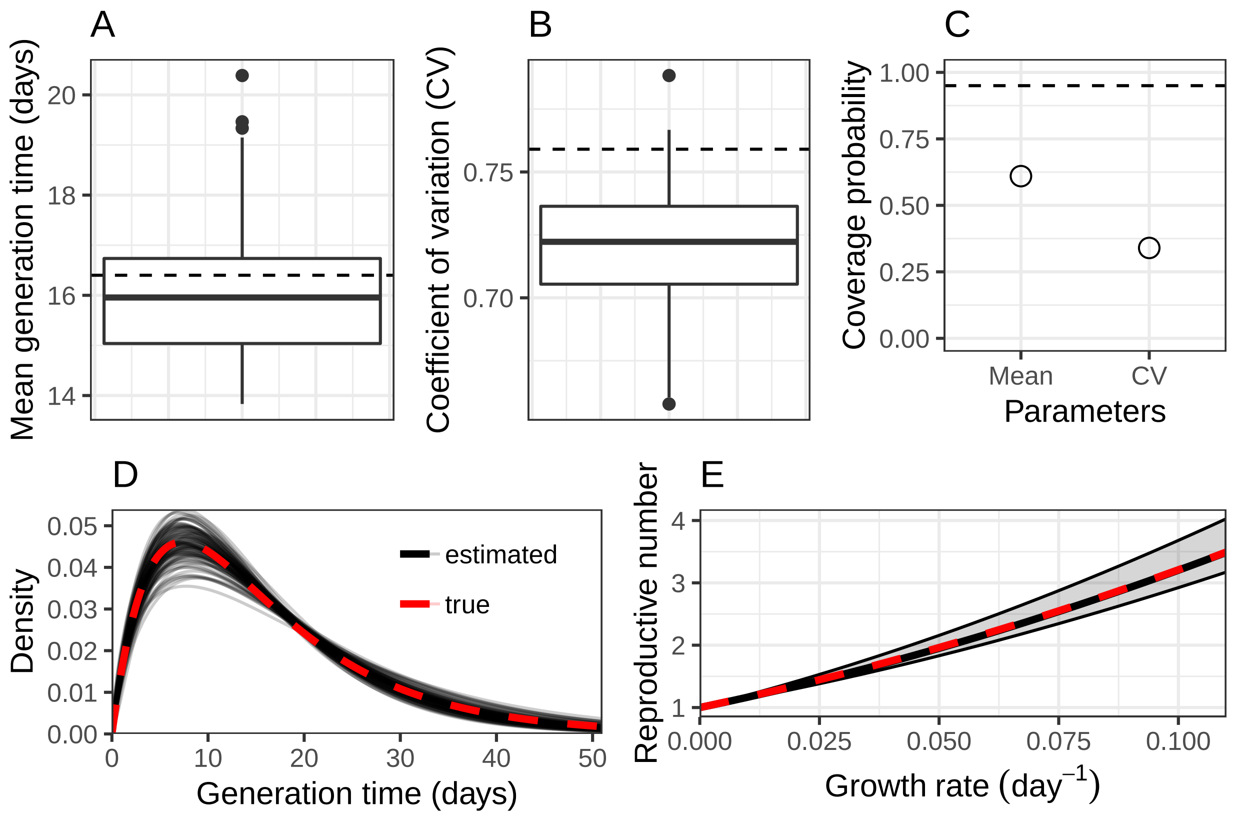


Figure A.3: **Wrong distributional assumption may result in biased estimates of the parameters of a generation-interval distribution.** We simulate a stochastic SEIR model on a homogenous network with $10^5$ individuals using Ebola-like parameters (1): mean latent period $1/\sigma = 5$ days, mean infectious period $1/\gamma = 11.4$ days, and the basic reproductive number $\mathcal{R}_0 = 2$. Then, we apply the individual-based method based on the first 1000 infections. (A-B) A boxplot of 100 estimates of the mean generation intervals and their coefficient of variations (CV). Dashed horizontal lines represent the true value. (C) Coverage probability of the mean generation intervals and their CVs based on the 95% confidence interval. Dashed horizontal lines represent the 95% coverage probability. (D) Estimated generation-interval distributions based on the gamma approximation and a true generation-interval distribution of an SEIR model. (E) Estimated $r$–$\mathcal{R}$ relationships based on the gamma approximation and a true $r$-$\mathcal{R}$ relationship of an SEIR model. Even though there are biases in the estimates of the parameters of the generation-interval distribution, we can still get an unbiased estimate of the $r$–$\mathcal{R}$ relationship: a shorter mean generation interval decreases $\mathcal{R}$ whereas a tighter generation-interval distribution increases $\mathcal{R}$ (2; 3).
